# Abcg2a is the functional homolog of human ABCG2 expressed at the zebrafish blood-brain barrier

**DOI:** 10.1101/2023.05.18.539313

**Authors:** Joanna R. Thomas, William J. E. Frye, Robert W. Robey, Andrew C. Warner, Donna Butcher, Jennifer L. Matta, Tamara C. Morgan, Elijah F. Edmondson, Paula B. Salazar, Suresh V. Ambudkar, Michael M. Gottesman

## Abstract

**Background:** A principal protective component of the mammalian blood-brain barrier (BBB) is the high expression of the multidrug efflux transporters P-glycoprotein (P-gp, encoded by *ABCB1*) and ABCG2 (encoded by *ABCG2*) on the lumenal surface of endothelial cells. The zebrafish P-gp homolog Abcb4 is expressed at the BBB and phenocopies human P-gp. Comparatively little is known about the four zebrafish homologs of the human *ABCG2* gene: *abcg2a*, *abcg2b*, *abcg2c*, and *abcg2d*. Here we report the functional characterization and brain tissue distribution of zebrafish ABCG2 homologs.

**Methods:** To determine substrates of the transporters, we stably expressed each in HEK-293 cells and performed cytotoxicity and fluorescent efflux assays with known ABCG2 substrates. To assess the expression of transporter homologs, we used a combination of RNAscope *in situ* hybridization probes and immunohistochemistry to stain paraffin-embedded sections of adult and larval zebrafish.

**Results:** We found Abcg2a had the greatest substrate overlap with ABCG2, and Abcg2d appeared to be the least functionally similar. We identified *abcg2a* as the only homolog expressed at the adult and larval zebrafish BBB, based on its localization to claudin-5 positive brain vasculature.

**Conclusions:** These results demonstrate the conserved function of zebrafish Abcg2a and suggest that zebrafish may be an appropriate model organism for the studying the role of ABCG2 at the BBB.

## Introduction

The blood-brain barrier (BBB) refers to the specialized protective adaptations of the brain microvasculature, posing a significant impediment to the delivery of therapeutics to the brain (1). Brain endothelial cells play a prominent role at the BBB, as they form tight junctions that prevent paracellular transport, thus constituting a physical barrier to compounds. They also express ATP-binding cassette (ABC) transporters that act as a chemical barrier by actively transporting compounds back into the blood supply, therefore limiting brain penetration (2). P-glycoprotein (P-gp, encoded by the *ABCB1* gene) and ABCG2 (encoded by the *ABCG2* gene) are the most highly expressed ABC transporters at the mammalian BBB. Knockout mouse models have consistently demonstrated the important role played by P-gp and ABCG2 at the BBB in limiting the brain penetration of therapeutics (3). For example, eight hours after oral administration of 10 mg/kg of the janus kinase (JAK) 1 and 2 inhibitor momelotinib, the brain concentration was 3.1-fold, 6.5-fold, and 48.4-fold higher in mice deficient in Abcb1a/b, Abcg2 or Abcb1a/b;Abcg2 compared to wild-type mice (4). Several targeted therapies are known to be restricted from the brain due to ABC transporter overexpression as demonstrated by knockout mouse models (5). However, mouse models have certain limitations.

Given its small size and amenability to higher throughput assays, the zebrafish has been suggested as a suitable *in vivo* model of the BBB. However, little is known about the role of ABC transporters at the zebrafish BBB. Zebrafish express 2 homologs of human *ABCB1*, termed *abcb4* and *abcb5* (6). While early studies with the anti-P-gp antibody C219 suggested that a zebrafish homolog of P-gp was expressed in brain capillaries (7, 8), the epitope for the C219 antibody is present in both zebrafish homologs of P-gp (8). We subsequently showed that Abcb4 is the sole P-gp homolog expressed at the zebrafish BBB and found that the substrate specificity of Abcb4 completely overlaps with that of P-gp, while Abcb5 confers resistance to a slightly narrower range of tested substrates (9). Much less work has been done to characterize the zebrafish homologs of *ABCG2*.

Zebrafish have 4 direct homologs of the human *ABCG2* gene—*abcg2a*, *abcg2b*, *abcg2c*, and *abcg2d*—yet relatively little is known about the function or expression of the transporters (10). Tsinkalovsky and colleagues hypothesized that an *ABCG2*-related gene was expressed in cells extracted from zebrafish kidney marrow, which serves a similar function as the bone marrow in mammals, based on the discovery of a side population of cells that accumulated low levels of Hoechst 33342, a fluorescent ABCG2 and P-gp substrate (11). This side population could be reduced by the addition of reserpine or fumitremorgin C (FTC), both of which are known to inhibit ABCG2-mediated transport (11). These observations were extended by Kobayashi et al., who isolated side population cells and found expression of both *abcg2a* and *abcg2c* (12). To determine which was responsible for transport of Hoechst 33342, they transfected cells with either gene and found that *abcg2a* overexpression resulted in decreased Hoechst 33342 retention (12). Additionally, expression of *abcg2a* and *abcg2b* was found to be high in the intestine, while *abcg2c* expression was high in the head kidney, body kidney, spleen, intestine, and gills, and *abcg2d* was not detected in any of the tissues examined (12). Given the paucity of data, we determined that a thorough investigation of the zebrafish homologs of human ABCG2 was needed. We hypothesized that an ABCG2 homolog is present at the zebrafish BBB. We therefore sought to determine which ABCG2 homologs are expressed at the zebrafish BBB and compare the substrate specificity of the various homologs to that of human ABCG2.

## Materials and Methods

### Chemicals

Mitoxantrone was obtained from Sigma-Aldrich (St. Louis, MO). Pheophorbide a was from Frontier Specialty Chemicals (Logan, UT). MLN-7243 and MLN-4924 were purchased from ChemieTek (Indianapolis, IN). THZ531 and Ko143 were obtained from Cayman Chemical (AnnArbor, MI). PF-3758309 and Gedatolisib were from ApexBio Technology (Houston, TX). CUDC-101, KS176 and tariquidar were from MedChem Express (Monmouth Junction, NJ). Elacridar was from Selleck chemicals (Houston, TX).

### Antibodies

A custom rabbit polyclonal anti-Abcg2a antibody was generated and purified by ProSci Incorporated (ProSci, Poway, CA). Rabbits were immunized with the immunogenic peptide MADTRVELTGVLDLQNHVN-C which corresponds to amino acids 1-19 in zebrafish Abcg2a (UniProt accession number Q2Q447). Commercially available rabbit monoclonal anti-beta actin (Cell Signaling Technologies, Danvers, MA; cat #4970, clone 13E5) was used for immunoblotting, while mouse monoclonal anti-claudin 5 (Invitrogen, Waltham, MA, cat #35-2500, clone 4C3C2) and mouse monoclonal anti-P-gp (Signet Laboratories, Dedham, MA, clone C219) were used for immunohistochemistry.

### Zebrafish housing and husbandry

Zebrafish (*Danio rerio*) wild-type strain TAB5 were housed with a 14-hour/10-hour light/dark cycle with a water temperature of 28.5°C in a recirculating aquatic system. Zebrafish husbandry and embryonic staging was conducted as previously described (13). Zebrafish embryos were raised in an incubator at 28.5°C in Instant Ocean (Blacksburg, VA) (60 mg/L DI water) and were euthanized at 7 days post-fertilization. All animal studies were compliant with the National Cancer Institute-Bethesda Animal Care and Use Committee approved study protocol (LCB-033-2).

### Cell lines

HEK-293 cells (ATCC, Manassas, VA) were transfected with empty pcDNA3.1 vector or with vector containing full-length *abcg2a* (NM_001042775), *abcg2b* (NM_001039066)*, abcg2c* (XM_005156523) or *abcg2d* (NM_001042772) (all from Genscript, Piscataway, NJ), flanked by a C-terminal FLAG tag sequence. The R-5 cell line was generated as previously described from HEK-293 cells transfected with full-length *ABCG2* (14). Transfected cells were cultured at 37°C and at 5% CO_2_ in EMEM (Quality Biological, Gaithersburg, MD) supplemented with 10% fetal calf serum, penicillin/streptomycin (Quality Biological) and 2 mg/ml G418 (Mediatech, Manassas, VA).

### Immunoblot

Total cell lysates were extracted from transfected HEK-293 cells using RIPA buffer and incubated at 37°C for 20 minutes. Lysates (30 μg protein) were resolved by gel electrophoresis on a NuPAGE™ 4 – 12% Bis-Tris gel (Invitrogen, Waltham, MA) and then transferred to a nitrocellulose membrane. The membrane was blocked in 5% non-fat milk in PBST (0.1% Tween 20) for 1 hour at room temperature, then probed overnight with our custom rabbit polyclonal anti-Abcg2a antibody (diluted 1:5000) and rabbit monoclonal anti-beta actin (Cell Signaling Technologies, Danvers, MA; cat #4970, clone 13E5, diluted 1:2000) while rocking at 4°C. The primary antibody was detected by incubating the membrane with IRDye® 680RD goat anti-rabbit IgG Secondary Antibody (LI-COR, Lincoln, NE), and measuring the fluorescence using the LiCor ODYSSEY CLx membrane scanner.

### Flow cytometry

Flow cytometry efflux assays with fluorescent human ABCG2 substrates were based on those previously described (15). Briefly, trypsinized cells were plated and incubated with fluorescent substrates (5 µM pheophorbide a or 5 µM mitoxantrone) in the presence or absence of ABCG2 inhibitors (10 µM KS176, 10 µM Ko143, 10 µM elacridar, 10 µM tariquidar) at 37°C for 30 minutes. The medium was removed, and cells were incubated in media in the presence or absence of inhibitors for 1 h. Cells were then washed and re-suspended in ice-cold PBS prior to analysis using a FACSCanto II flow cytometer (BD Biosciences San Jose, CA) and data were then analyzed with FlowJo software (v 10.8.1, Tree Star, Inc, Ashland, OR). Pheophorbide a and mitoxantrone fluorescence was detected with a 635-nm red diode laser and a 670-nm filter. At least 10,000 events were collected for each sample. Samples incubated with fluorescent substrates without inhibitors are defined as the ‘efflux’ histogram, and unstained cell samples are the control cell autofluorescence histogram.

### Cytotoxicity assays

Trypsinized cells were plated at 5,000 cells/well in white opaque 96-well plates and allowed to adhere overnight. Drugs were then added, with each concentration tested in triplicate, and cells were incubated for 72 hours. Cell growth was assessed using the CellTiter-Glo (Promega, Madison, WI) reagent, according to the manufacturer’s instructions. Luminescence was measured on a Tecan Infinite M200 Pro microplate reader (Tecan Group, Morrisville, NC). 50% growth inhibition (GI_50_) values were calculated at the drug concentration at which 50% luminescence was observed in comparison to untreated cells. Relative resistance (RR) values are the ratio of the GI_50_ values of the transporter transfected versus the empty vector-transfected HEK-293 cells.

### Immunofluorescence and RNAScope probes

Adult zebrafish were euthanized by immersion in a lethal dose of Tricane Methanesulfonate for 30 minutes, prior to fixation in 10% neutral buffered formalin (NBF) for 24 hours at room temperature. Larvae were anesthetized using Tricane Methanesulfonate and euthanized via rapid chilling (hypothermic shock), prior to fixation in 10% NBF for 24 hours at room temperature. Zebrafish were incubated in 0.5 M EDTA (pH 8) at room temperature for 7 days with gentle agitation. Before embedding and processing, zebrafish were washed twice with nuclease-free PBS. Paraffin-embedded samples were processed as previously described (9). Briefly, coronal and transverse sections were cut from the paraffin block at 5 µm and placed on positively charged slides for RNA in situ hybridization (ISH) (RNAscope) and immunohistochemistry (IHC).

The expression of *Danio rerio abcg2a*, *abcg2b*, *abcg2c*, *abcg2d*, and *abcb4* genes were detected by staining 5 µm FFPE sections with the RNAscope^®^ 2.5 LS Probe-Dr-abcg2a (Advanced Cell Diagnostics (ACD) (Newark, CA), Cat# 493571), Dr-abcg2b-C2 (ACD, Cat# 516878-C2), Dr-abcg2c-C2 (ACD, Cat# 516888-C2), Dr-abcg2d-C2 (ACD, Cat# 516898-C2), or Dr-abcb4-C2 (ACD, Cat# 493558-C2) with the RNAscope LS Multiplex Fluorescent Assay (ACD, Cat# 322800) using the Bond RX auto-stainer (Leica Biosystems, Deer Park, IL) with a tissue pretreatment of 15 minutes at 95°C with Bond Epitope Retrieval Solution 2 (Leica Biosystems), 15 minutes of Protease III (ACD) at 40°C and 1:750 dilution of OPAL570 or OPAL690 reagents (Akoya Biosciences, Marlborough, MA). Immunohistochemistry for claudin-5 co-localization was performed after RNAScope staining, using an anti-claudin-5 antibody (Invitrogen, Waltham, MA, cat #35-2500) at a 1:50 dilution, the custom anti-Abcg2a antibody (ProSci, Poway, CA) at a 1:10,000 dilution, or an anti-P-gp antibody (Signet Laboratories, Dedham, MA, clone C219) at a 1:200 dilution, for 30 minutes using the Bond Polymer Refine Kit (Leica Biosystems, cat# DS9800) minus the DAB and Hematoxylin with a 1:150 dilution of OPAL520 reagent for 10 min. The RNAscope 3-plex LS Multiplex Negative Control Probe (Bacillus subtilis dihydrodipicolinate reductase (dapB) gene) in channels C1, C2, and C3, ACD, Cat# 320878) followed by IHC with no primary antibody was used as an ISH and IHC negative control. Slides were digitally imaged using an Aperio ScanScope FL Scanner (Leica Biosystems).

### Homology modeling

The Modeller package, integrated with the PyMOL Molecular Graphics System (version 2.5.2) in its Pymol 3.0 plugin, was used to generate three-dimensional structural models for zebrafish Abcg2a-d paralogs. As a template, the cryo-EM structures of human ABCG2 in complex with topotecan (PDB: 7NEZ) and mitoxantrone (PDB: 6VXI) were used (16, 17). Alignment of predicted Abcg2a-d structures to the cryo-EM structure of ABCG2 was performed using the European Bioinformatics Institute (EMBL-EBI) Clustal Omega program (https://www.ebi.ac.uk/Tools/msa/clustalo/).

The degree of the sequence homology was color coded corresponding to EMBL-EBI Clustal Multiple Sequence Alignment consensus symbols. Fully conserved residues (EBI symbol * (asterisk)) are colored black; conserved residues with strongly similar properties (EBI symbol: (colon)), approximately equivalent to scoring > 0.5 in the Gonnet PAM 250 matrix, are colored green; residues with weakly similar properties (EBI symbol. (period)), approximately equivalent to scoring =< 0.5 and > 0 in the Gonnet PAM 250 matrix are colored yellow, and non-conservative substitutions are depicted with red color.

## Results

### Characterization of HEK-293 cells stably expressing zebrafish Abcg2a-d

Human embryonic kidney (HEK-293) cells were stably transfected to express an empty vector, or a vector containing human *ABCG2*, zebrafish *abcg2a*, *abcg2b*, *abcg2c*, or *abcg2d*. Zebrafish gene constructs included a 5’ FLAG tag sequence to enable protein immunodetection, as there are no commercially available antibodies that cross-react with any zebrafish ABCG2 homologs. Positive FLAG staining was observed in Abcg2a-d-expressing cell pellets (Fig. 1A). RNAscope probe positive signal was observed only in the corresponding transfected cell line, demonstrating specific detection of *abcg2a-d* homologs (Fig. 1B). The level of RNAscope probe staining was comparable among samples, indicating a similar level of mRNA expression among the corresponding cell lines. The typical punctate staining pattern seen in RNAscope probe images is not observed here due to the high signal density.

**Fig. 1.**
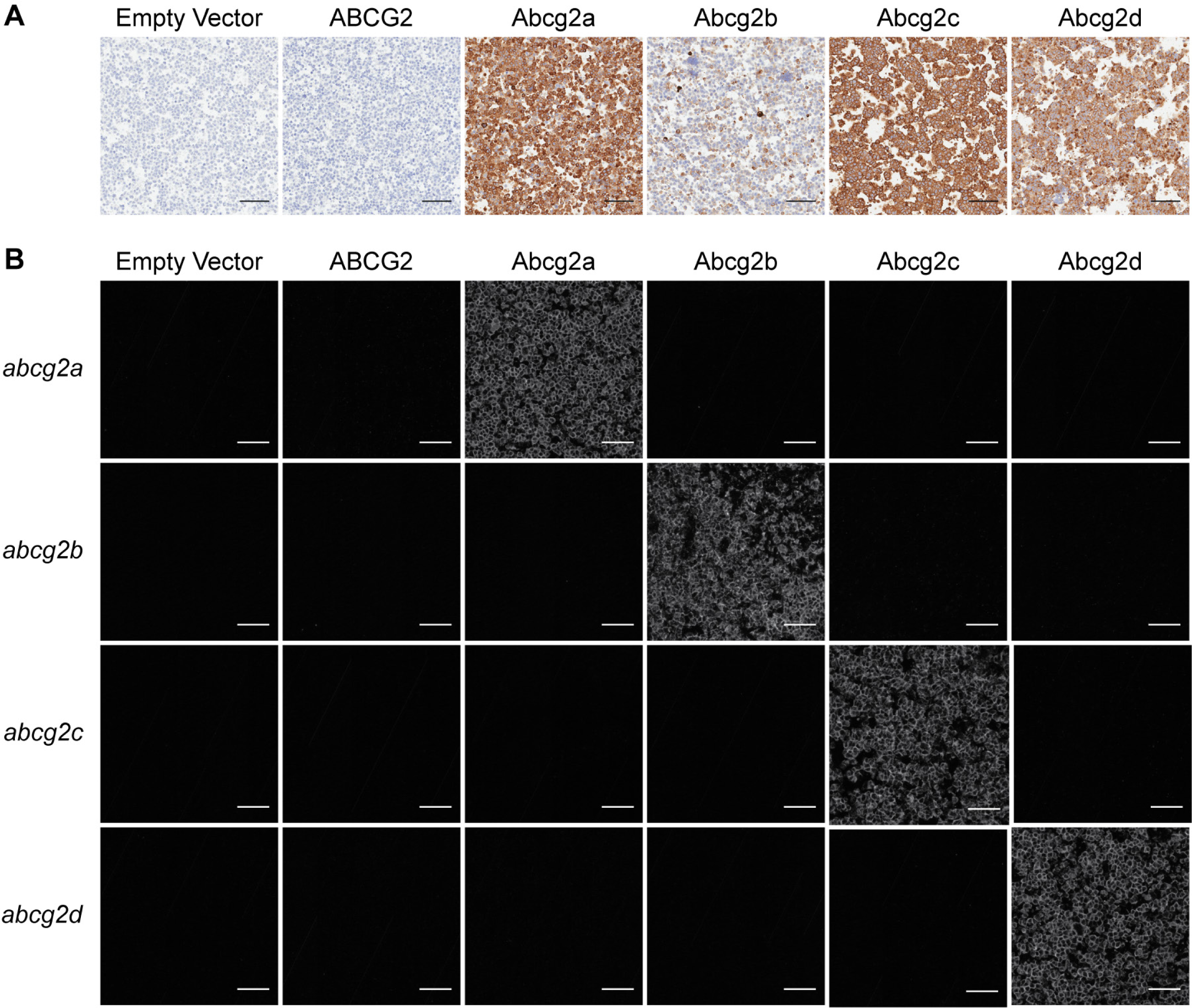
Characterization of HEK-293 cells transfected to express an empty vector, human ABCG2, zebrafish Abcg2a, Abcg2b, Abcg2c, or Abcg2d. Paraffin-embedded cells were probed with (A) an anti-FLAG tag antibody, or (B) RNAscope probes to detect *abcg2a*, *abcg2b*, *abcg2c*, or *abcg2d* mRNA. Scale bar = 100 μM.

### Zebrafish Abcg2 homologs confer resistance to ABCG2 substrates

To determine if zebrafish Abcg2a-d were able to confer resistance to known cytotoxic substrates of ABCG2, cytotoxicity assays were performed (18–23) and quantified by the drug concentration that produces 50% cell growth inhibition (GI_50_). Among the zebrafish Abcg2 homologs, Abcg2a conferred the highest level of resistance to MLN7243 (a ubiquitin activating enzyme inhibitor), MLN4924 (NEDD8 activating enzyme inhibitor), and CUDC-101 (histone deacetylase and receptor tyrosine kinase inhibitor) (Fig. 2, Table S1). Abcg2d conferred the least resistance to THZ 531 (CDK12/13 inhibitor), CUDC-101, and gedatolisib (an inhibitor of phosphatidylinositol-3-kinase (PI3K) and mammalian target of rapamycin (mTOR)). All zebrafish Abcg2 homologs conferred resistance to PF-3758309 (p21-activated kinase 4 inhibitor). Abcg2b and Abcg2c often conferred similar levels of resistance to the same substates, as evidenced by results with MLN7243, MLN4924, THZ 531, CUDC-101 and gedatolisib.

**Fig. 2.**
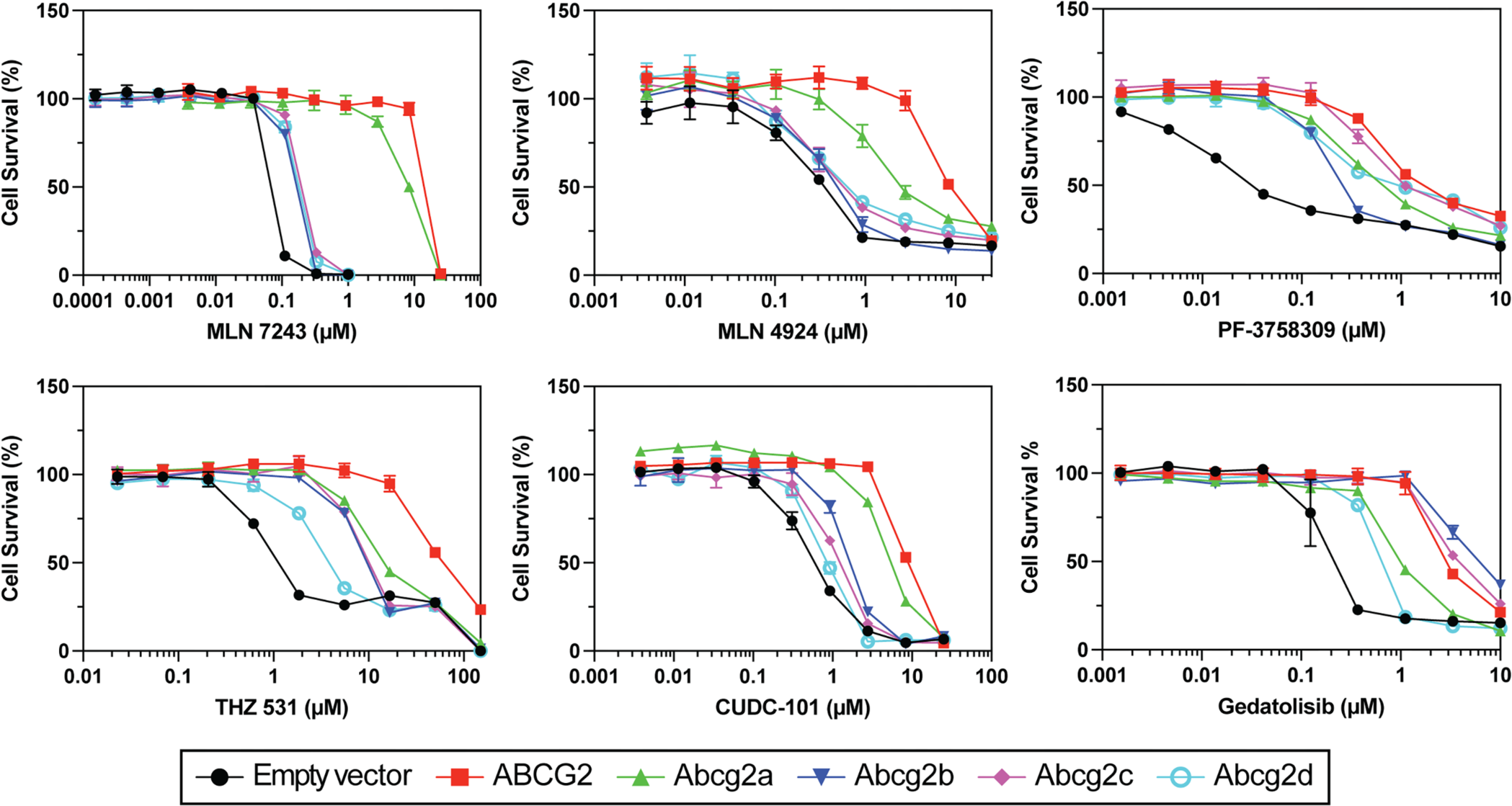
Zebrafish ABCG2 homologs confer resistance to cytotoxic ABCG2 substrates. HEK-293 cells stably expressing empty vector, ABCG2 or Abcg2a-d were treated with known cytotoxic ABCG2 substrates for 3 days. GI_50_ and relative resistance values are summarized in Table S1.

### Zebrafish Abcg2 homologs can efflux fluorescent ABCG2 substrates

We assessed the ability of Abcg2a-d homolog-expressing cells to efflux the fluorescent ABCG2 substrates mitoxantrone and pheophorbide a. Cells were incubated with 5 µM mitoxantrone or pheophorbide a in the absence or presence of 10 µM of the ABCG2 inhibitors KS176, Ko143, elacridar, or tariquidar, and flow cytometry was used to measure intracellular fluorescence. When co-incubated with inhibitors, ABCG2-expressing cells exhibit an increase in intracellular fluorescence evidenced by a shift to the right of histograms corresponding to the inhibitors compared to uninhibited efflux peaks, with little overlap between the histograms (Fig. 3). There was a rightward inhibited peak shift for Abcg2a with mitoxantrone, and to a greater extent with pheophorbide a, indicating Abcg2a can efflux both substrates. Pheophorbide a also appears to be a substrate for Abcg2b and Abcg2c. Neither mitoxantrone nor pheophorbide a were transported by Abcg2d, indicated by the overlay of the efflux and inhibited histograms. In addition to substrate specificities, these data also show all 4 ABCG2 inhibitors were able to inhibit Abcg2a, Abcg2b, and Abg2c.

**Fig. 3.**
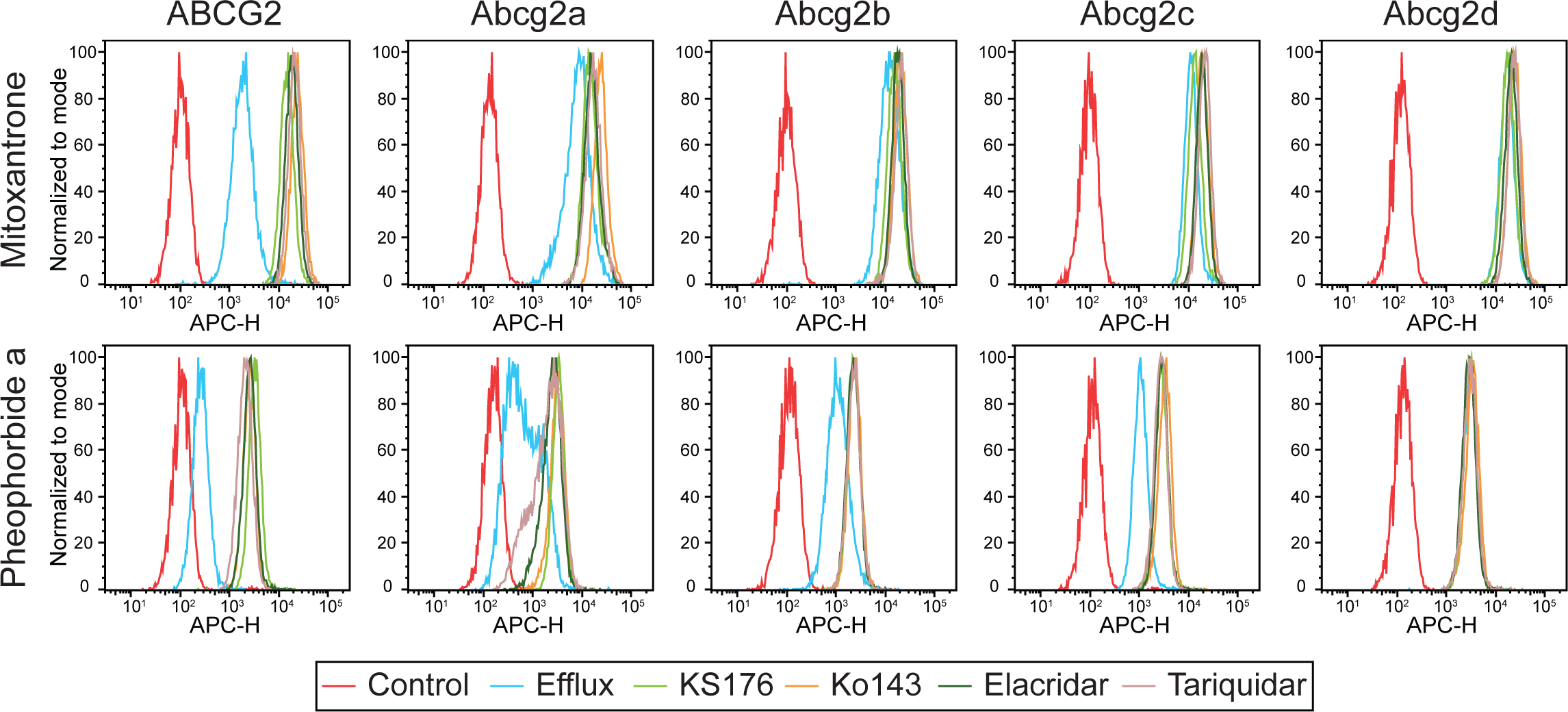
Zebrafish ABCG2 homologs differentially efflux fluorescent ABCG2 substrates. HEK293 cells overexpressing empty vector (EV), ABCG2 or Abcg2a-d were treated with the fluorescent ABCG2 substrates mitoxantrone and pheophorbide a in the absence or presence of the ABCG2 inhibitors KS176, Ko143, elacridar, and tariquidar. Representative histograms are shown from at least 3 biological replicates of intracellular fluorescence measured by flow cytometry. The control condition (red) of untreated cells shows cell autofluorescence, and the efflux condition (blue) indicates cells treated with substrate without inhibitor.

### Abcg2a-d homology modelling

Despite the fact Abcg2d is the homolog with second highest amino acid identity to human ABCG2 (Table S2), it has the least substrate overlap. To seek insights into differences in substrate specificity between ABCG2 and Abcg2a-d paralogs, we employed homology modeling techniques. We aligned predicted 3-D structures of Abcg2a-d to cryo-EM structures of ABCG2 bound to the substrate drugs topotecan or mitoxantrone (Fig. S1), focusing on the putative drug binding pocket (Fig. S2), and color-coded amino acid residues based on their level of conservation. As expected, based on total amino acid identity and our functional data, the drug binding pocket of Abcg2a exhibited the highest conservation to ABCG2. In contrast, the drug binding pocket of Abcg2b and Abcg2c displayed the greatest differences and Abcg2d presented an intermediate level of conservation.

### *abcg2a* is expressed at the adult zebrafish blood-brain barrier

Since ABCG2 is highly expressed at the mammalian BBB, we hypothesized that one or more of the ABCG2 homologs would localize to the adult zebrafish brain. Positive staining for *abcg2a* RNA was observed in adult brain sections, evidenced by a punctate signal which localized to the vasculature (Fig. 4). No positive staining was observed for *abcg2b* or *abcg2c* in the brain. Low levels of *abcg2d* staining were observed in the brain parenchyma, but the transporter was not localized to the vasculature. No differences were observed between brain regions or between males and females.

**Fig. 4.**
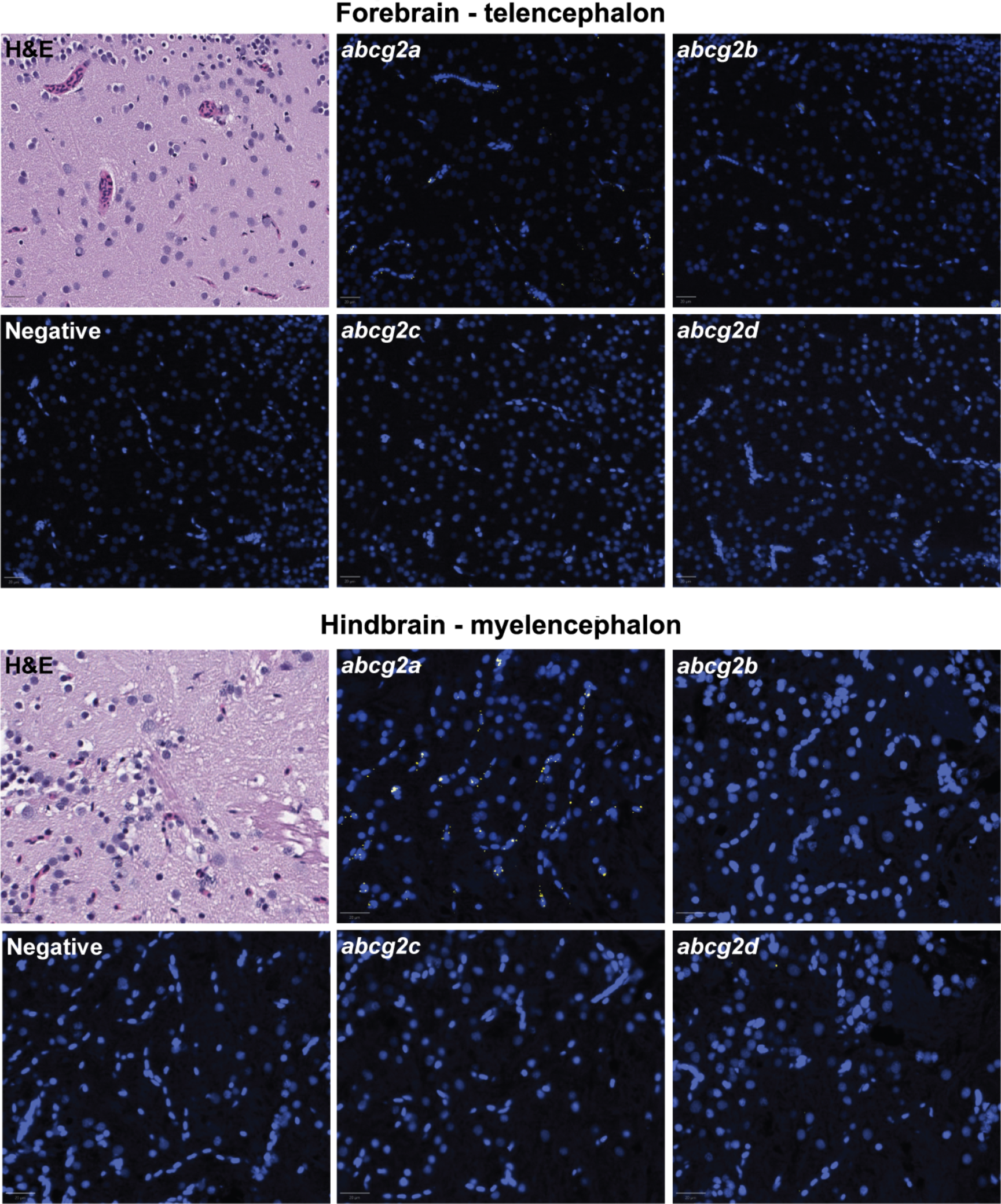
*Abcg2a* is the only ABCG2 homolog detected in adult zebrafish brain vasculature. Paraffin-embedded adult zebrafish sections were stained with hematoxylin and eosin (H&E) probed with RNAscope probes (yellow) to detect *abcg2a*, *abcg2b*, *abcg2c*, or *abcg2d* RNA, and DAPI (blue). Each punctate dot signal represents amplification of a single target RNA molecule. Sections shown are representative cross sections of the telencephalon and myelencephalon. Scale bar = 20 μm.

To determine if the vasculature where *abcg2a* localized had BBB properties, we co-stained with the endothelial tight junction protein claudin-5, a canonical marker of the BBB, using an antibody that cross-reacts with mammalian and zebrafish claudin-5 homologs (24). The positive signal for *abcg2a* co-localized with claudin-5 positive vasculature, indicating it is expressed at the adult zebrafish BBB (Fig 5). The zebrafish P-gp homolog *abcb4*, which we previously determined is expressed at the zebrafish BBB (9), also localized to the same claudin-5 positive vessels as *abcg2a*.

**Fig. 5.**
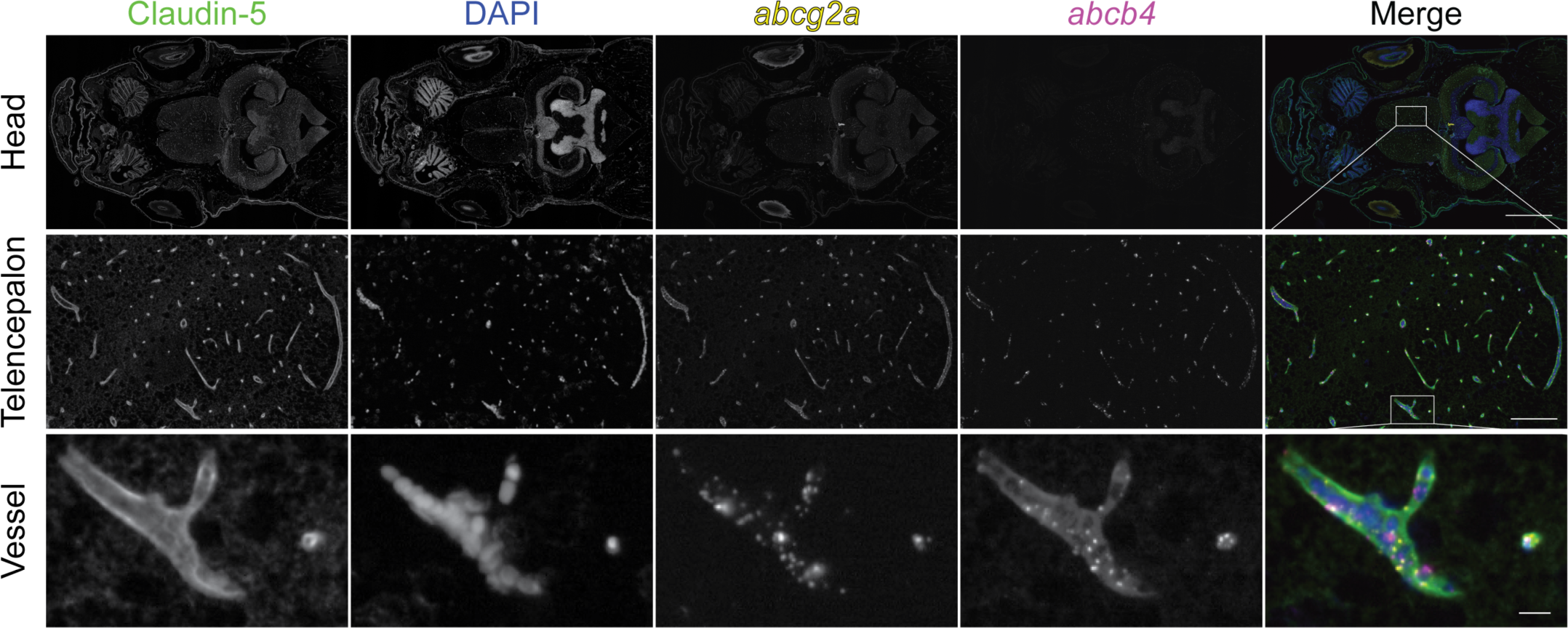
*abcg2a* and *abcb4* are expressed in claudin-5 positive adult brain vasculature. Paraffin-embedded adult zebrafish sections were probed with RNAscope probes to detect *abcg2a* (yellow) and *abcb4* (magenta) mRNA, an antibody against claudin-5 (green), and DAPI (blue). Head scale bar = 1 mm, telencephalon scale bar = 100 μm, vessel scale bar = 10 μm.

### Abcg2a and Abcb4 are co-expressed at the blood-brain barrier

There are no commercially available antibodies that cross-react with Abcg2a, so we generated a custom antibody and validated its specificity (Fig. S1). The commercial antibody C219, which recognizes human and murine P-gp, cross-reacts with the zebrafish P-gp homologs Abcb4 and Abcb5. We previously confirmed using RNAscope probes that the positive C219 signal in the brain vasculature corresponds to Abcb4 expression (9). In sequential adult brain sections, we observed positive staining with Abcg2a and C219 antibodies in claudin-5 positive vasculature throughout the brain. This confirms the co-expression of Abcg2a and Abcb4 protein at the adult zebrafish BBB.

### *abcg2a* is expressed at the developing larval blood-brain barrier

The zebrafish BBB begins to form between 1- and 3-days post fertilization (dpf) and becomes progressively restrictive until 10 dpf (8, 25). We therefore sought to determine if any *abcg2* paralogs localized to the developing larval BBB. At 7 dpf we detected *abcg2a* localized to claudin-5 positive brain vasculature (Fig. 6). At 5 dpf *abcg2a* and *abcb4* both localize to claudin-5 positive brain vasculature (Fig. S2B). In concordance with our adult staining, *abcg2b, abcg2c* and *abcg2d* do not localize to the larval brain vasculature. Sex is not differentiated in 7 dpf larvae and therefore cannot be investigated as a variable.

**Fig. 6.**
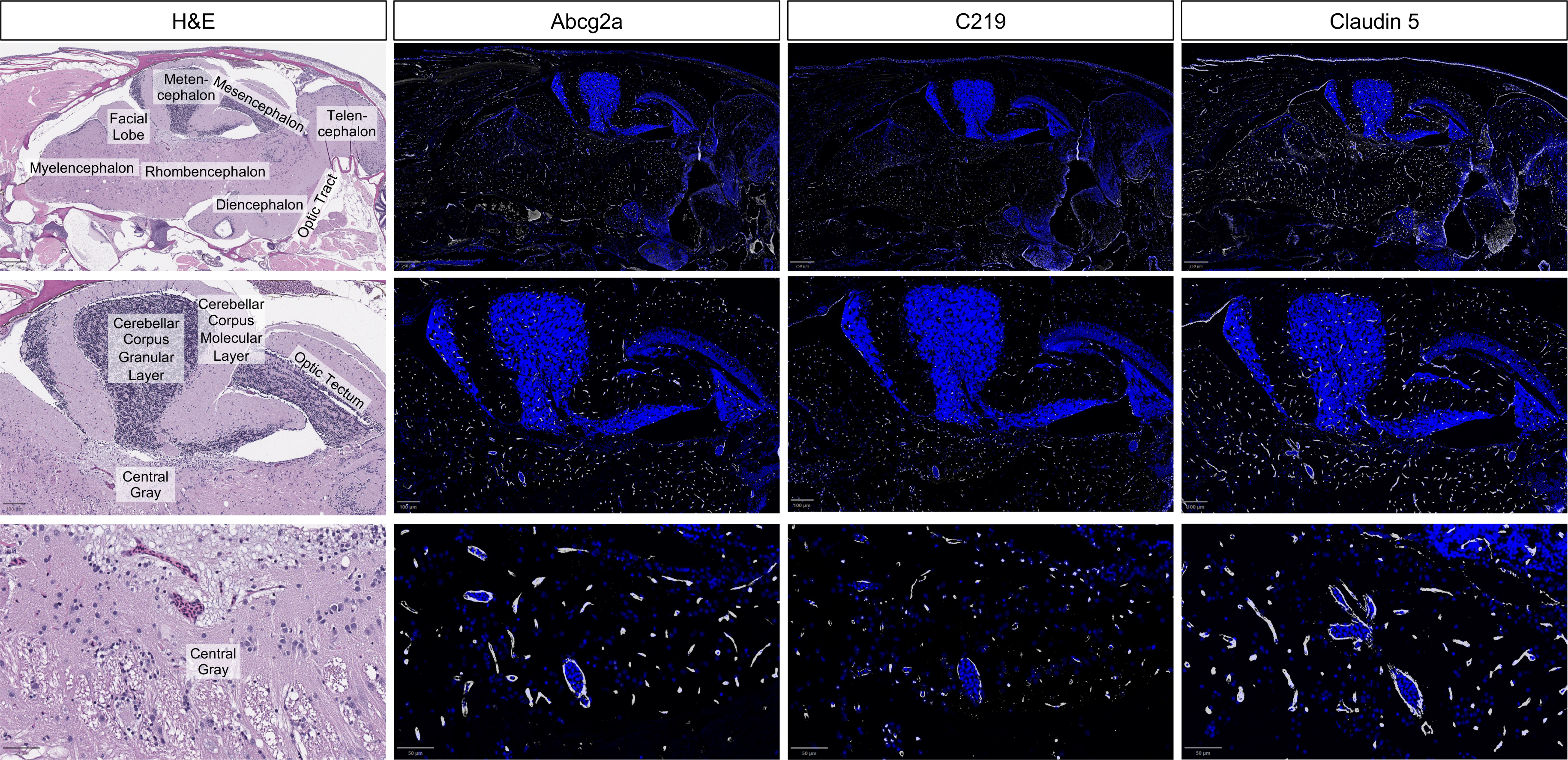
Abcg2a and Abcb4 protein localize to claudin-5 positive vasculature. Sagittal section of an adult zebrafish brain stained with hematoxylin and eosin (H&E) and probed with anti-Abcg2a, C219 (cross-reacts with Abcb4), and claudin-5 antibodies. Descending row panels show the entire brain (upper), section of the brain (middle), and an enlarged region (lower). Antibody staining is pseudo colored in gray, and DAPI in blue. Scale bar upper = 250 μm, middle = 100 μm, lower = 50 μm.

**Fig. 7.**
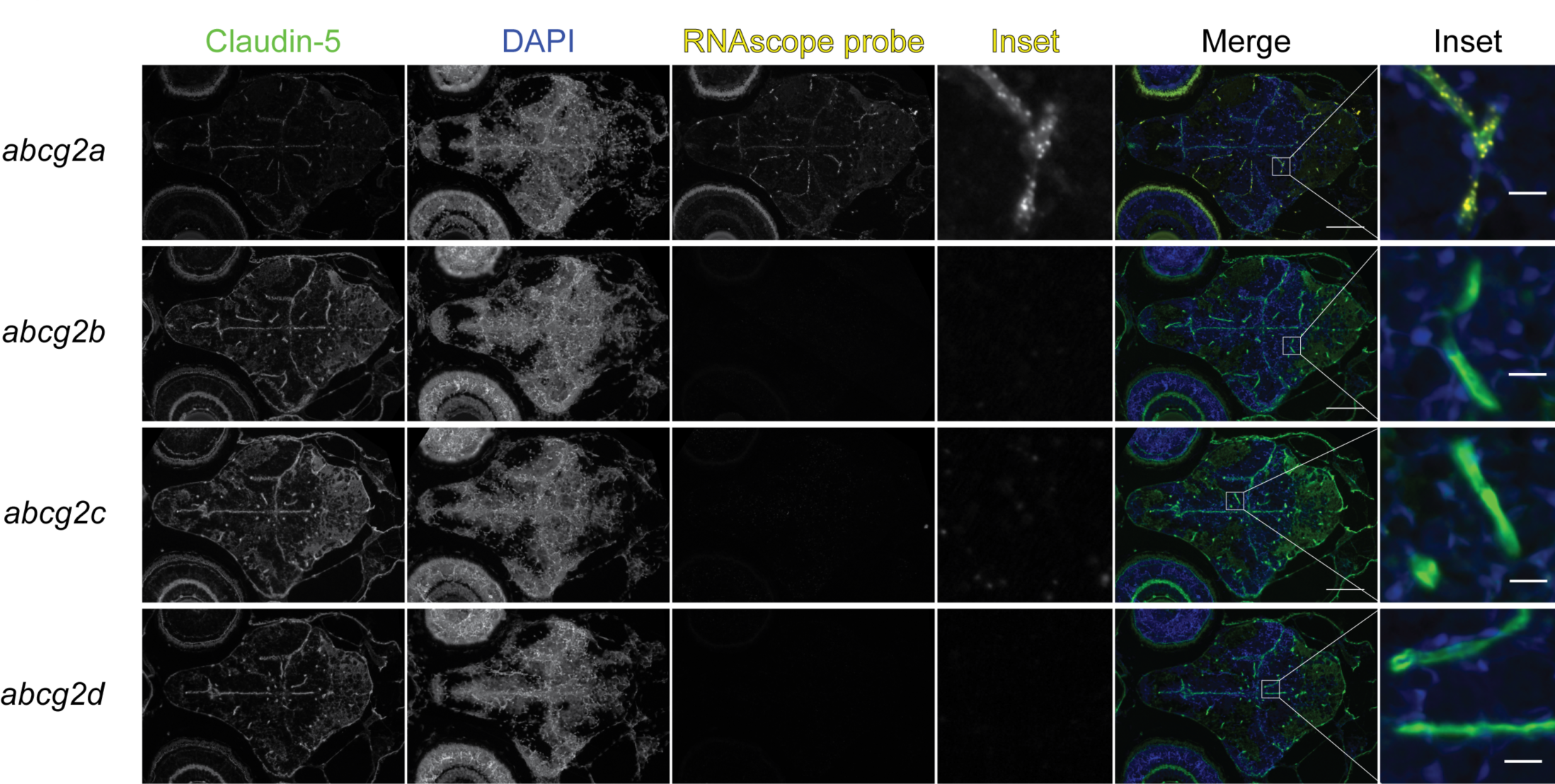
*abcg2a* is expressed in claudin-5 positive larval brain vasculature. Paraffin-embedded 7 dpf larval zebrafish sections were probed with RNAscope probes (yellow) to detect *abcg2a* mRNA, an antibody against claudin-5 (green) and DAPI (blue). Scale bar = 100 μm, inset scale bar = 10 μm.

## Discussion

Here we report our characterization of the function and brain distribution of the zebrafish homologs of ABCG2. We used stably-transfected HEK-293 cells as a model to study the function of each of the zebrafish Abcg2 homologs in isolation, and a panel of 8 substrates and 4 inhibitors of ABCG2 in cytotoxicity and fluorescent efflux assays. We found that Abcg2a had the greatest substrate overlap with ABCG2 and was able to transport all assayed ABCG2 substrates. Abcg2d appeared to be the most divergent from ABCG2 and had the lowest GI_50_ for 4 out of the 6 cytotoxic substrates and did not efflux either fluorescent substrate. We expected Abcg2a to be the most similar to ABCG2, given the fact it has the highest amino acid homology (61.4%, Table S2), effluxes Hoechst 33342, as does ABCG2, and is inhibited by verapamil which also inhibits ABCG2 (12). It was surprising to us that Abcg2d appeared to be the least functionally similar to ABCG2, as it has the second highest amino acid homology (57.4%) to ABCG2 and was predicted based on synteny to be the most direct homolog of ABCG2 (10). Our homology modelling did demonstrate that Abcg2d had a more divergent substrate binding pocket than Abcg2a compared to ABCG2. These variations in the structure of the drug binding pocket could help elucidate the differences in substrate specificity observed among Abcg2 homologs. Thus, we found that all 4 zebrafish homologs were able to transport select ABCG2 substrates to varying degrees and that the substrate profile of Abcg2a is most similar to human ABCG2.

Overall, the ABCG2 substrates we assayed seem to be weaker substrates for the zebrafish homologs, evidenced by the lower GI_50_ values in cytotoxicity assays compared to ABCG2 (Fig 2, Table S1), and the less pronounced peak shift between the uninhibited and inhibited conditions in fluorescent efflux assays (Fig 3). The only exception to this was in the case of gedatolisib, which was transported slightly more efficiently by Abcg2b (GI_50_ 4.06±1.46 μM) than ABCG2 (GI_50_ 3.84±1.53μM). The ABCG2 inhibitors KS176, Ko143, elacridar, and tariquidar were all able to inhibit Abcg2a, -b, and -c, at the same concentration used with ABCG2 (10 μM). It is unknown if they inhibit Abcg2d, and this will need to be assessed with additional fluorescent substrates, or in cytotoxicity assays with identified Abcg2d substrates. Some human P-gp inhibitors are less effective against the zebrafish homologs Abcb4 and Abcb5 in the case of select substrates. For example, the P-gp (and ABCG2) inhibitor elacridar was less effective than the P-gp inhibitor valspodar at inhibiting Abcb4 and Abcb5 rhodamine 123 transport (9). Our data suggest that all 4 inhibitors are similarly effective at inhibiting the Abcg2a-c transport of pheophorbide a, evidenced by the overlay of histograms corresponding to inhibitors (Fig. 3). A wider range of ABCG2 substrates and varying inhibitor concentrations are needed to assess substrate-specific inhibitor efficacy for zebrafish Abcg2 paralogs. One reasonable hypothesis is that the zebrafish Abcg2 paralogs have evolved to recognize toxic materials in the marine environment, and/or zebrafish-specific metabolites and therefore their drug specificities differ somewhat from human ABCG2.

Expression of zebrafish ABC transporters in mammalian cells is a tractable method to assess their function in isolation; however, it also comes with the caveat that zebrafish proteins may not function in these heterologous expression systems as they do *in vivo* in zebrafish. Fluorescent ABCG2 substrates could be administered *in vivo* to zebrafish larvae in the presence or absence of inhibitors in conjunction with fluorescent microscopy to confirm if they are Abcg2 substrates, as has been done for zebrafish Abcb4 with rhodamine 123 and calcein-AM (6).

Zebrafish *abcg2a* was the only homolog expressed at the BBB in adult and larval zebrafish. Abcg2a and Abcb4, expressed in claudin-5 positive vasculature in the adult brain, likely represent the relevant ABCG2 and P-gp homologs at the zebrafish BBB. This suggests they may function in concert to protect the brain parenchyma, as they do in mammals. In humans, an analysis of transporter density by brain region (normalized by vascular density) showed P-gp density is 9% lower in gray versus white matter, and is lower in the parietal cortex (19%), temporal cortex (15%) and hippocampus (17%) relative to the thalamus; and ABCG2 did not vary by brain region but was 33% higher in gray versus white matter (26). We did not observe any differences in the distribution of Abcb4 or Abcg2a in gray versus white matter or by brain region, although quantification may reveal differences indistinguishable by a pathologist’s eye. One of the major species differences between the human and rodent BBB is the higher expression of both P-gp and ABCG2 homologs in rodents (27), resulting in a higher brain to plasma ratio of P-gp substrates in humans (28). Quantification of transporter levels at the zebrafish BBB has not yet been reported. Also, the subcellular localization of Abcb4 and Abcg2a has not been determined, but the increase of brain to vascular ratio of Abcb4 substrates upon transporter inhibition and knockout is consistent with luminal expression (29, 30).

It will be interesting to determine at what developmental stage *abcg2a* and *abcb4* localize to the BBB, as this will inform the utility of larval zebrafish for BBB studies. We detected *abcg2a* and *abcb4* at the BBB of 5 dpf larvae, although the expression of both has been reported from 24 hours post fertilization (hpf) in a BBB vasculature subcluster (hema.28 vasculature blood-brain barrier) from whole-embryo single-cell sequencing (31, 32). At 5 dpf, the most mature age represented in the dataset of Sur and colleagues, *abcg2a* and *abcb4* are the first and second most highly expressed ABC transporters in the BBB subcluster, indicating they may be the predominant, but not the only, ABC transporters expressed.

In conclusion, we have shown that zebrafish ABCG2 homologs are functionally conserved, with Abcg2a appearing to be most similar to ABCG2. The localization of Abcg2a to the BBB and its substrate and inhibitor overlap with ABCG2 indicate that the zebrafish has potential utility for the study of Abcg2 activity at the BBB. Our data, and the literature, demonstrate functional similarities between Abcg2a and ABCG2, and Abcb4 and P-gp, giving credence to the idea that the zebrafish could be a relevant model system to assess the brain penetration of substrate drugs (6, 9, 12).

## Abbreviations

ABC: ATP-binding cassette
BBB: blood-brain barrier; dpf, days post-fertilization
ABCG2: human protein
*ABCG2*: human gene
Abcg2a-d: zebrafish proteins
*abcg2a-d*: zebrafish genes
ISH: *in situ hybridization*
IHC: immunohistochemistry
hpf: hours post-fertilization.

## Acknowledgements

We are grateful for the editorial assistance of George Leiman and William Rhodes.

## Authors’ Contributions

Conception and design of work: JRT, RWR, MMG

Acquisition, analysis, or interpretation of data: JRT, WJEF, RWR, ACW, DB, JLM, TCM, EFE, PBS

Preparation and review of manuscript: JRT, WJEF, RWR, ACW, EFE, PBS, SVA, MMG

All authors read and approved the final manuscript.

## Availability of data

The datasets supporting the conclusions of this article are included within the article and its additional files.

## Declarations

### Ethics approval

Zebrafish studies were performed under a protocol approved by the National Cancer Institute Animal Care and Use Committee (IACUC Number LCB-033).

## References

1. O’Brown NM, Pfau SJ, Gu C. Bridging barriers: a comparative look at the blood-brain barrier across organisms. Genes Dev. 2018;32(7-8):466–78.

2. Schulz JA, Hartz AMS, Bauer B. ABCB1 and ABCG2 Regulation at the Blood-Brain Barrier: Potential New Targets to Improve Brain Drug Delivery. Pharmacol Rev. 2023;75(5):815–53.

3. Agarwal S, Hartz AM, Elmquist WF, Bauer B. Breast cancer resistance protein and P-glycoprotein in brain cancer: two gatekeepers team up. Curr Pharm Des. 2011;17(26):2793–802.

4. Durmus S, Xu N, Sparidans RW, Wagenaar E, Beijnen JH, Schinkel AH. P-glycoprotein (MDR1/ABCB1) and breast cancer resistance protein (BCRP/ABCG2) restrict brain accumulation of the JAK1/2 inhibitor, CYT387. Pharmacol Res. 2013;76:9–16.

5. Lagas JS, Vlaming ML, Schinkel AH. Pharmacokinetic assessment of multiple ATP-binding cassette transporters: the power of combination knockout mice. Mol Interv. 2009;9(3):136–45.

6. Fischer S, Kluver N, Burkhardt-Medicke K, Pietsch M, Schmidt AM, Wellner P, et al. Abcb4 acts as multixenobiotic transporter and active barrier against chemical uptake in zebrafish (Danio rerio) embryos. BMC Biol. 2013;11:69.

7. Umans RA, Taylor MR. Zebrafish as a model to study drug transporters at the blood-brain barrier. Clin Pharmacol Ther. 2012;92(5):567–70.

8. Fleming A, Diekmann H, Goldsmith P. Functional characterisation of the maturation of the blood-brain barrier in larval zebrafish. PLoS One. 2013;8(10):e77548.

9. Robey RW, Robinson AN, Ali-Rahmani F, Huff LM, Lusvarghi S, Vahedi S, et al. Characterization and tissue localization of zebrafish homologs of the human ABCB1 multidrug transporter. Sci Rep. 2021;11(1):24150.

10. Annilo T, Chen ZQ, Shulenin S, Costantino J, Thomas L, Lou H, et al. Evolution of the vertebrate ABC gene family: analysis of gene birth and death. Genomics. 2006;88(1):1–11.

11. Tsinkalovsky O, Vik-Mo AO, Ferreira S, Laerum OD, Fjose A. Zebrafish kidney marrow contains ABCG2-dependent side population cells exhibiting hematopoietic stem cell properties. Differentiation. 2007;75(3):175–83.

12. Kobayashi I, Saito K, Moritomo T, Araki K, Takizawa F, Nakanishi T. Characterization and localization of side population (SP) cells in zebrafish kidney hematopoietic tissue. Blood. 2008;111(3):1131–7.

13. Westerfield M. The Zebrafish Book: A Guide for the Laboratory Use of Zebrafish (Danio Rerio): University of Oregon Press; 2000.

14. Robey RW, Honjo Y, Morisaki K, Nadjem TA, Runge S, Risbood M, et al. Mutations at amino acid 482 in the ABCG2 gene affect substrate and antagonist specificity. Br J Cancer. 2003;89:1971–8.

15. Robey RW, Steadman K, Polgar O, Morisaki K, Blayney M, Mistry P, Bates SE. Pheophorbide a is a specific probe for ABCG2 function and inhibition. Cancer Res. 2004;64(4):1242–6.

16. Kowal J, Ni D, Jackson SM, Manolaridis I, Stahlberg H, Locher KP. Structural Basis of Drug Recognition by the Multidrug Transporter ABCG2. J Mol Biol. 2021;433(13):166980.

17. Orlando BJ, Liao M. ABCG2 transports anticancer drugs via a closed-to-open switch. Nat Commun. 2020;11(1):2264.

18. Wu ZX, Yang Y, Wang JQ, Narayanan S, Lei ZN, Teng QX, et al. Overexpression of ABCG2 Confers Resistance to MLN7243, a Ubiquitin-Activating Enzyme (UAE) Inhibitor. Front Cell Dev Biol. 2021;9:697927.

19. Liu C, Xing W, Yu H, Zhang W, Si T. ABCB1 and ABCG2 restricts the efficacy of gedatolisib (PF-05212384), a PI3K inhibitor in colorectal cancer cells. Cancer Cell Int. 2021;21(1):108.

20. Wu CP, Hsiao SH, Su CY, Luo SY, Li YQ, Huang YH, et al. Human ATP-Binding Cassette transporters ABCB1 and ABCG2 confer resistance to CUDC-101, a multi-acting inhibitor of histone deacetylase, epidermal growth factor receptor and human epidermal growth factor receptor 2. Biochem Pharmacol. 2014;92(4):567–76.

21. Bradshaw-Pierce EL, Pitts TM, Tan AC, McPhillips K, West M, Gustafson DL, et al. Tumor P-Glycoprotein Correlates with Efficacy of PF-3758309 in in vitro and in vivo Models of Colorectal Cancer. Front Pharmacol. 2013;4:22.

22. Gao Y, Zhang T, Terai H, Ficarro SB, Kwiatkowski N, Hao MF, et al. Overcoming Resistance to the THZ Series of Covalent Transcriptional CDK Inhibitors. Cell Chem Biol. 2018;25(2):135–42 e5.

23. Wei LY, Wu ZX, Yang Y, Zhao M, Ma XY, Li JS, et al. Overexpression of ABCG2 confers resistance to pevonedistat, an NAE inhibitor. Exp Cell Res. 2020;388(2):111858.

24. Abdelilah-Seyfried S. Claudin-5a in developing zebrafish brain barriers: another brick in the wall. Bioessays. 2010;32(9):768–76.

25. Umans RA, Henson HE, Mu F, Parupalli C, Ju B, Peters JL, et al. CNS angiogenesis and barriergenesis occur simultaneously. Dev Biol. 2017;425(2):101–8.

26. Kannan P, Schain M, Kretzschmar WW, Weidner L, Mitsios N, Gulyas B, et al. An automated method measures variability in P-glycoprotein and ABCG2 densities across brain regions and brain matter. J Cereb Blood Flow Metab. 2017;37(6):2062–75.

27. Uchida Y, Ohtsuki S, Katsukura Y, Ikeda C, Suzuki T, Kamiie J, Terasaki T. Quantitative targeted absolute proteomics of human blood-brain barrier transporters and receptors. J Neurochem. 2011;117(2):333–45.

28. Syvänen S, Lindhe O, Palner M, Kornum BR, Rahman O, Långström B, et al. Species differences in blood-brain barrier transport of three positron emission tomography radioligands with emphasis on P-glycoprotein transport. Drug Metab Dispos. 2009;37(3):635–43.

29. Park J, Kim H, Alabdalla L, Mishra S, McHaourab H. Generation and characterization of a zebrafish knockout model of abcb4, a homolog of the human multidrug efflux transporter P-glycoprotein. Hum Genomics. 2023;17(1):84.

30. Kim SS, Im SH, Yang JY, Lee YR, Kim GR, Chae JS, et al. Zebrafish as a Screening Model for Testing the Permeability of Blood-Brain Barrier to Small Molecules. Zebrafish. 2017;14(4):322–30.

31. Sur A, Wang Y, Capar P, Margolin G, Farrell JA. Daniocell: National Institute of Child Health and Human Development; 2023 [updated 2023/03/21. v1.0:[Available from: https://daniocell.nichd.nih.gov/.

32. Sur A, Wang Y, Capar P, Margolin G, Prochaska MK, Farrell JA. Single-cell analysis of shared signatures and transcriptional diversity during zebrafish development. Dev Cell. 2023.

