## Supplemental Figures for "Abcg2a is the functional homolog of human ABCG2 expressed at the zebrafish blood-brain barrier"

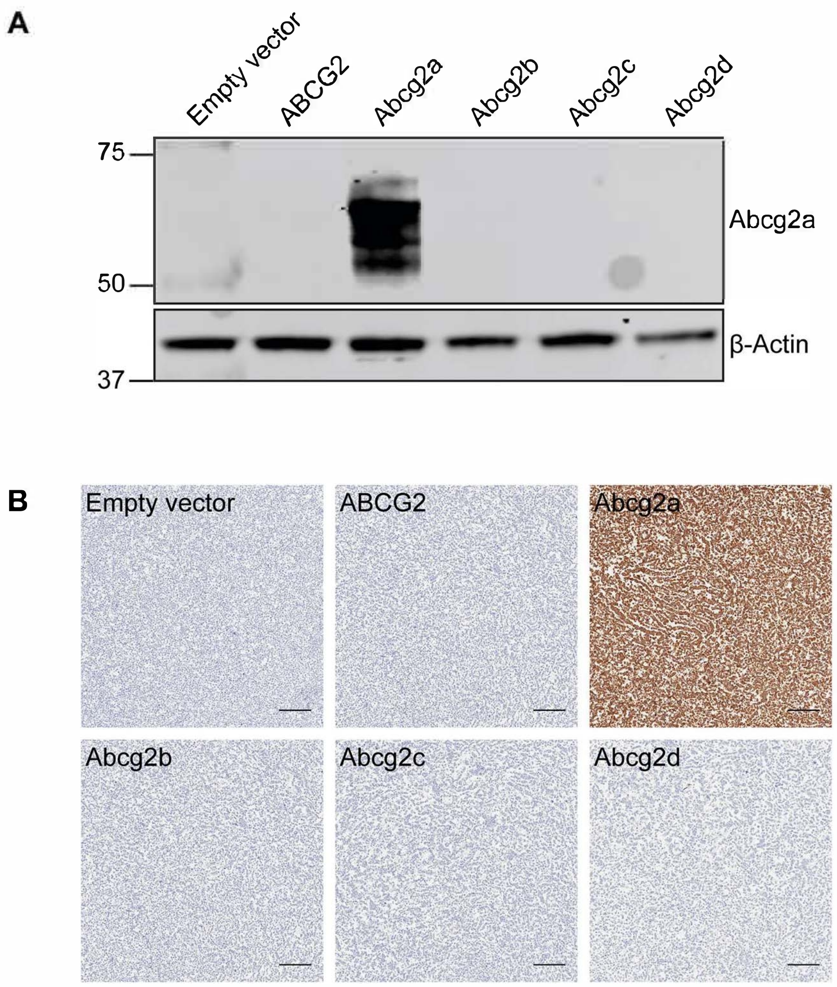


**Fig. S1.** Abcg2a antibody validation. (A) Immunoblot of total cell lysates and (B) immunohistochemistry of pellets of transfected HEK-293 cells expressing an empty vector, ABCG2, Abcg2a, Abcg2b, Abcg2c, or Abcg2d. Positive signal is only observed in Abcg2a-expressing cells. Scale bar = 100 µm.

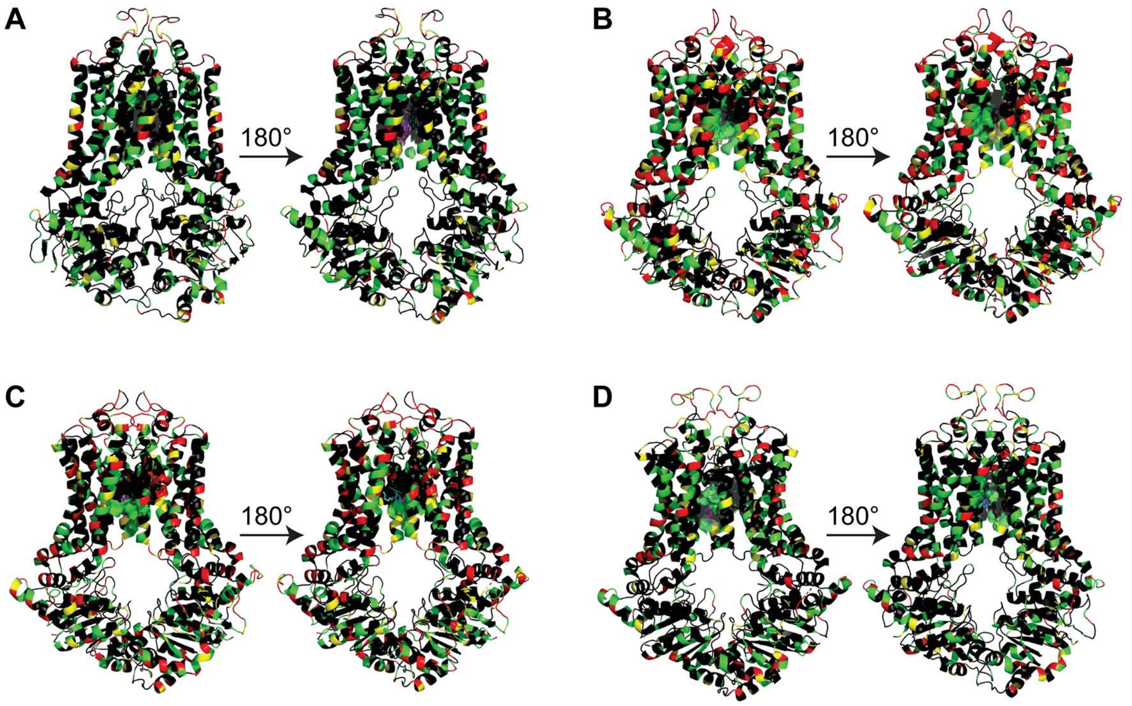


**Fig. S2.** 3-D homology modeling of sequence similarity of zebrafish (A) Abcg2a, (B) Abcg2b, (C) Abcg2c, and (D) Abcg2d. Predicted structures of zebrafish Abcg2a-d paralogs were aligned to the cryo-EM structure of human ABCG2 (PDB IDs: 7NEZ and 6VXI for topotecan and mitoxantrone bound structures, respectively). Structures are presented in cartoon mode with a 180Árotated view, and the predicted drug binding pocket is presented in surface mode. Amino acids are color coded black (fully conserved), green (conservative substitution), yellow (semi-conservative substitution), and red (non-conservative substitution) based on similarity to ABCG2. Clustal amino acid sequence comparisons comparing Abcg2 paralogs with ABCG2 calculated percent identities of 63.26%, 47.28%, 47.92%, and 61.31% for Abcg2a, Abcg2b, Abcg2c, and Abcg2d, respectively.

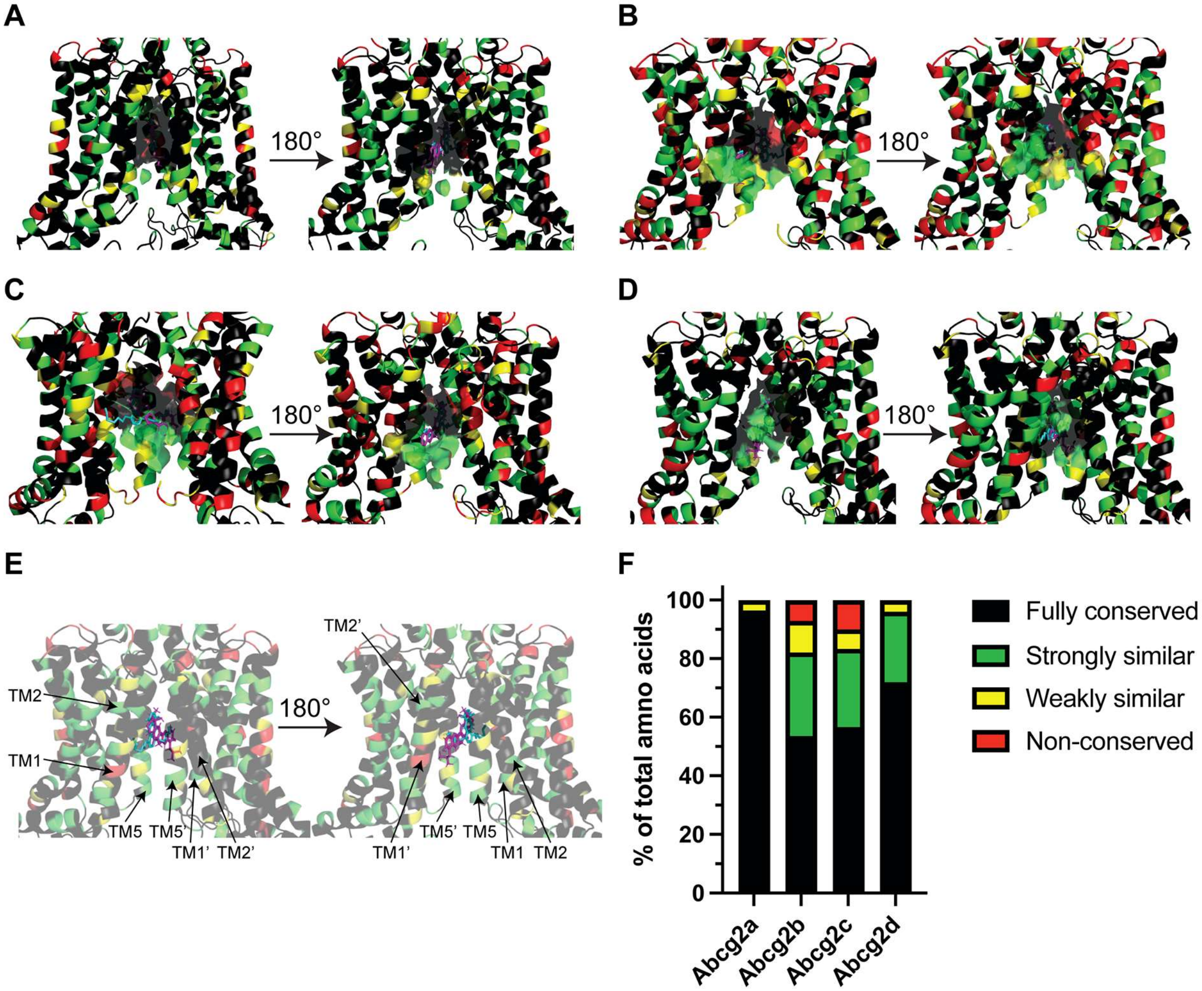


**Fig. S3.** 3-D homology modeling of amino acid similarity in the substrate binding pocket of (A) Abcg2a, (B) Abcg2b, (C) Abcg2c, and (D) Abcg2d. The transmembrane region is presented in cartoon mode with a 180° rotated view. Predicted drug-binding pockets, shown in surface mode, focused on residues within a 4.5 Å proximity to the ligands topotecan (magenta) and mitoxantrone (cyan) as identified in human ABCG2 (PDB IDs: 7NEZ and 6VXI, respectively). (E) A transparent view of panel (A) with the transmembrane helices labelled from each monomer in the homodimer. (F) Quantification of the percent of total amino acids in the predicted binding pocket that are fully conserved (black), conservative substitutions (green), semi-conservative substitutions (yellow), or non-conservative substitutions (red), based on similarity to ABCG2. Color coding in the structures (A-E) is consistent with the graph (F) and Fig. S2.

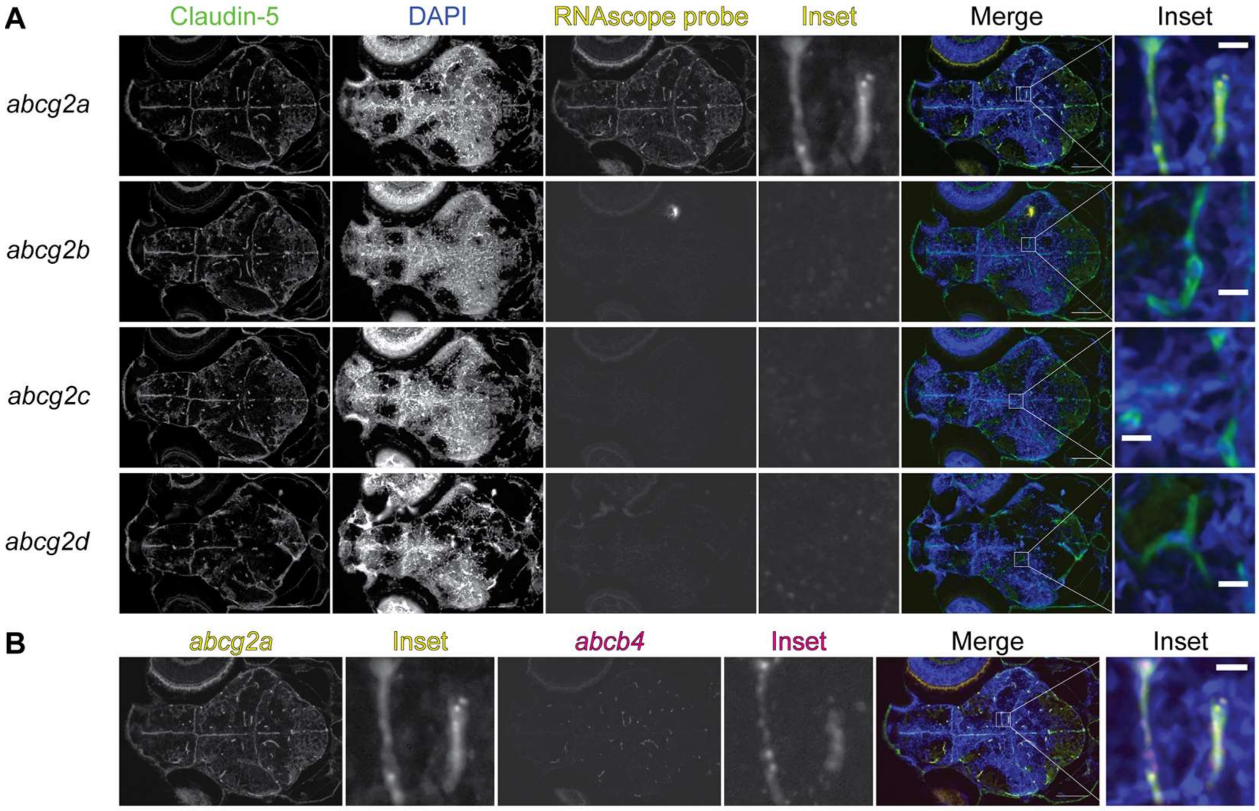


**Fig. S4.** *abcg2a* and *abcb4* are expressed in claudin-5 positive 5 dpf larval brain vasculature. (A) Paraffin-embedded 5 dpf larval zebrafish sections were probed with RNAscope probes (yellow) to detect *abcg2a-d* mRNA, an antibody against claudin-5 (green) and DAPI (blue). (B) Co-staining of the *abcg2a* probed section with an *abcb4* RNAscope probe. Scale bar = 100 εm, inset scale bar = 10 εm.

**Table S1:** Cross-resistance profile of known ABCG2 cytotoxic substrate drugs with zebrafish Abcg2a-d[a]

| Compound | $GI_{50}$ Empty vector (µM) | $GI_{50}$ ABCG2 (µM) | RR* ABCG2 | $GI_{50}$ Abcg2a (µM) | RR* Abcg2a | $GI_{50}$ Abcg2b (µM) | RR* Abcg2b | $GI_{50}$ Abcg2c (µM) | RR* Abcg2c | $GI_{50}$ Abcg2d (µM) | RR* Abcg2d |
|---|---|---|---|---|---|---|---|---|---|---|---|
| MLN 7243 | 0.069±.0050 | 15.4±.22 | 223 | 7.49±.2.8 | 109 | 0.151±.051 | 2.19 | 0.200±.020 | 2.90 | 0.179±.023 | 2.60 |
| MLN 4924 | 0.420±.090 | 7.32±1.7 | 17.6 | 2.44±.17 | 5.86 | 0.520±.052 | 1.25 | 0.487±.040 | 0.430 | 0.353±.19 | 0.850 |
| PF-3758309 | 0.068±.030 | 2.36±.54 | 34.8 | 1.26±.53 | 18.5 | 0.413±.15 | 6.09 | 1.97±.71 | 29.1 | 1.54±.65 | 22.7 |
| THZ 531 | 1.29±.14 | 65.9±3.7 | 51.1 | 18.3±3.6 | 14.2 | 10.5±.1.4 | 8.32 | 11.4±1.0 | 8.80 | 3.99±.24 | 3.09 |
| CUDC-101 | 0.780±.10 | 11.3±2.0 | 14.5 | 5.42±1.4 | 6.96 | 2.09±.19 | 2.68 | 1.69±.29 | 2.17 | 1.04±.26 | 1.33 |
| Gedatolicib | 0.200±.010 | 3.84±1.5 | 19.1 | 1.39±.19 | 6.86 | 4.06±1.5 | 20.1 | 3.26±.49 | 16.2 | 0.820±.50 | 4.07 |

[a] All compounds were tested in at least 3 biological replicates. Results are mean $GI_{50}$ values with +/- standard deviation.

* Relative resistance (RR) value is the ratio of GI50 values of ABCG2, Abcg2a-d overexpressing cells, to the empty vector

**Table S2:** UniProt percent amino acid identity matrix for human and zebrafish ABCG2 homologs[a]

| | Abcg2c | Abcg2b | Abcg2d | Abcg2a | ABCG2 |
|---|---|---|---|---|---|
| Abcg2c | 100 | 63.05 | 47.23 | 46.23 | 47.61 |
| Abcg2b | 63.05 | 100 | 48.53 | 47.86 | 47.55 |
| Abcg2d | 47.23 | 48.53 | 100 | 65.61 | 57.41 |
| Abc2a | 46.23 | 47.86 | 65.61 | 100 | 61.43 |
| ABCG2 | 47.61 | 47.55 | 57.41 | 61.43 | 100 |

[a]Uniprot accession numbers: Zebrafish Abcg2c, Q08CU5; Abcg2b, Q2Q44; Abcg2d, Q2Q444. Human ABCG2, Q9UNQ0
